# Cameo: A Python Library for Computer Aided Metabolic Engineering and Optimization of Cell Factories

**DOI:** 10.1101/147199

**Authors:** João G. R. Cardoso, Kristian Jensen, Christian Lieven, Anne Sofie Lærke Hansen, Svetlana Galkina, Moritz Beber, Emre Özdemir, Markus J. Herrgård, Henning Redestig, Nikolaus Sonnenschein

## Abstract

Computational systems biology methods enable rational design of cell factories on a genomescale and thus accelerate the engineering of cells for the production of valuable chemicals and proteins. Unfortunately, for the majority of these methods’ implementations are either not published, rely on proprietary software, or do not provide documented interfaces, which has precluded their mainstream adoption in the field. In this work we present cameo, a platform-independent software that enables *in silico* design of cell factories and targets both experienced modelers as well as users new to the field. It is written in Python and implements state-of-the-art methods for enumerating and prioritizing knock-out, knock-in, over-expression, and down-regulation strategies and combinations thereof. Cameo is an open source software project and is freely available under the Apache License 2.0. A dedicated website including documentation, examples, and installation instructions can be found at http://cameo.bio. Users can also give cameo a try at http://try.cameo.bio.

## INTRODUCTION

The engineering of cells for the production of chemicals and proteins affects all areas of our modern lives. Beer, yogurt, flavoring, detergents, and insulin represent just a few products which are unimaginable without biotechnology. Engineered cells may further provide solutions to many of mankind’s greatest challenges like global climate, multiple drug resistance, and overpopulation, by producing fuels, novel antibiotics, and food from renewable feedstocks. Manipulating cells to perform tasks that they did not evolved for, however, is challenging and requires significant investments and personnel in order to reach economically viable production of target molecules (Lee and Kim, 2015).

A central task in developing biotechnological production processes is to reroute metabolic fluxes towards desired products in cells. This task is particularly prone to failure due to our limited understanding of the underlying biology and the complexity of the metabolic networks in even the simplest of organisms. In line with other recent technological advancements, like high-fidelity genome editing through CRISPR/Cas9 (Sander and Joung, 2014) and DNA synthesis costs dropping (Kosuri and Church, 2014), modeling methods are increasingly used to accelerate cell factory engineering, helping to reduce development time and cost (Meadows et al., 2016).

Genome-scale models of metabolism (GEMs) (McCloskey et al., 2013) are of particular interest in this context as they predict phenotypic consequences of genetic and environmental perturbations affecting cellular metabolism (O’Brien et al., 2015). These models have been developed throughout the past 15 years for the majority of potential cell factory host organisms ranging from bacteria to mammalian cells. A large repertoire of algorithms has been published that utilize GEMs to compute cell factory engineering strategies composed of over-expression, down-regulation, deletion, and addition of genes (see Maia et al., 2016; Machado and Herrgård, 2015). Unfortunately, most of these algorithms are not easily accessible to users as they have either been published without implementation (e.g. using pseudo code or mathematical equations to describe the method) or the implementation provided by the authors is undocumented or hard to install. These problems significantly limit the ability of metabolic engineers to utilize computational design tools as part of their workflow.

## RESULTS

Cameo is open source software written in Python that alleviates these problems and aims to make *in silico* cell factory design broadly accessible. On the one hand it enables cell factory engineers to enumerate and prioritize designs without having to be experts in metabolic modeling themselves. On the other hand it aims to become a comprehensive library of published methods by providing method developers with a library that simplifies the implementation of new cell factory design methods.

Cameo provides a high-level interface that can be used without knowing any metabolic modeling or how different algorithms are implemented (see Supplementary Notebook 8 [v0.10.3, current]). In fact, the most minimal form of input that cameo requires is simply the desired product, for example vanillin.

**from cameo import** api

api.design(product=‘vanillin’)

This function call will run the workflow depicted in Figure 1. It is also possible to call the same functionality from the command line. Firstly, it enumerates native and heterologous production pathways for a series of commonly used host organisms and carbon sources. Then it runs a whole suite of design algorithms available in cameo to generate a list of metabolic engineering strategies, which can then be ranked by different criteria (maximum theoretical yield, number of genetic modifications etc.).

**Figure 1.**
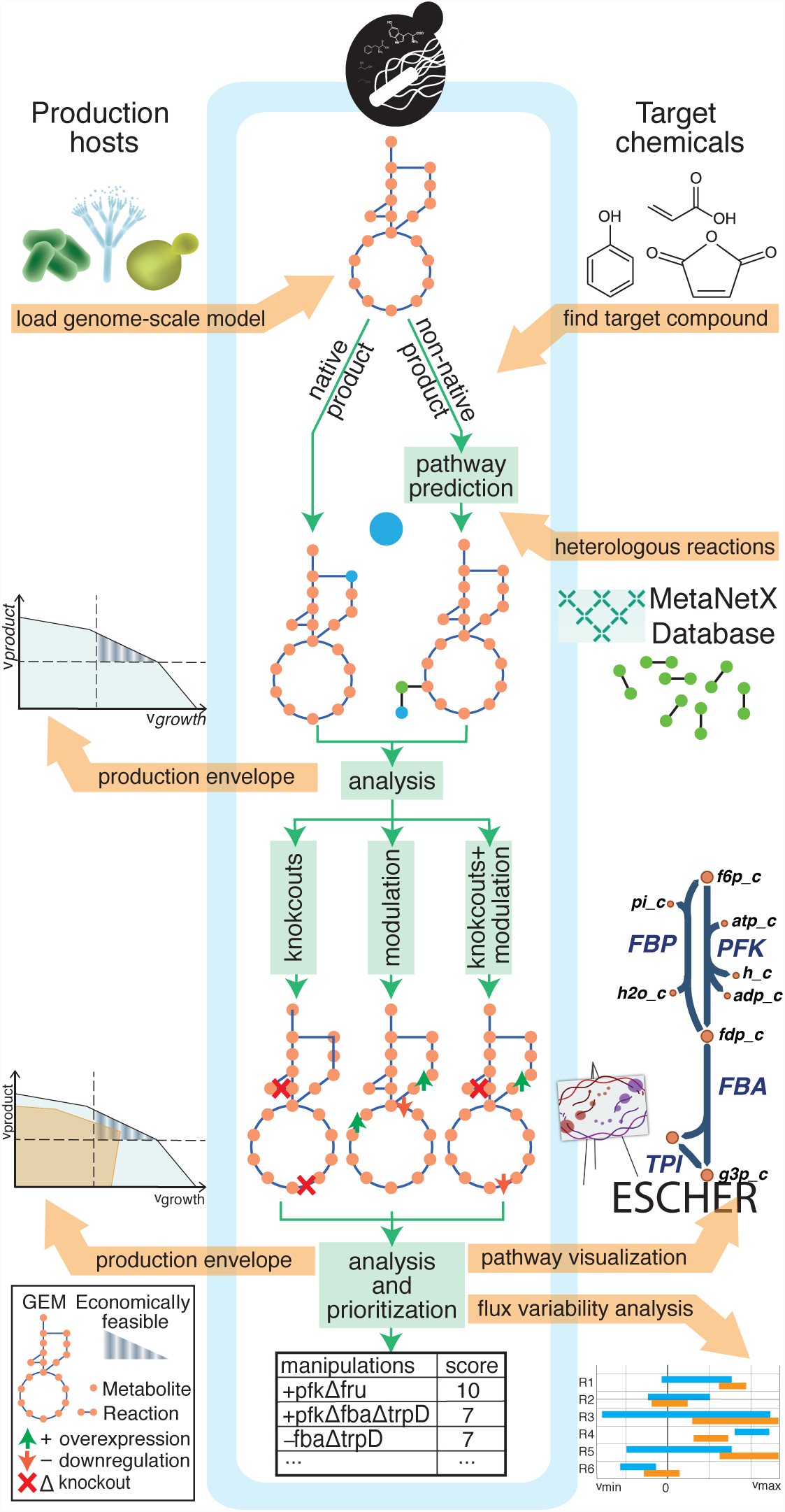
Cell factory design workflow with cameo. The first step is to import a metabolic model from a file or using a web service. Next, the user needs to select a target product. If the target product is a non-native chemical, shortest heterologous production pathways can be enumerated to determine a suitable route to the product (Pharkya et al., 2004). Potential production pathways can then be compared using production envelopes, i.e., visualizations of the trade-off between production rate and organism growth rate (see Supplementary Notebook 4 [v0.10.3, current]). After a production pathway has been chosen, a number of different design methods are used to compute the genetic modifications (designs) necessary to achieve the production goal (see Supplementary Notebooks 5 [v0.10.3, current] and 6 [v0.10.3, current]). In the end, the computed designs can be sorted using different criteria relevant to the actual implementation in the lab and economic considerations such as the number of genetic modifications needed and maximum theoretical product yield. Furthermore, a number of results can be further visualized using the pathway visualization tool Escher (King et al., 2015)

More advanced users can easily customize this workflow by providing models for other host organisms, changing parameters and algorithms, and of course by including their own methods.

In order to become a community project and attract further developers, cameo has been developed as a modular Python package that has been extensively documented and tested using modern software engineering practices like test-driven development and continuous integration/deployment on travis-ci.org (Figure 2 shows an overview of the package organization).

**Figure 2.**
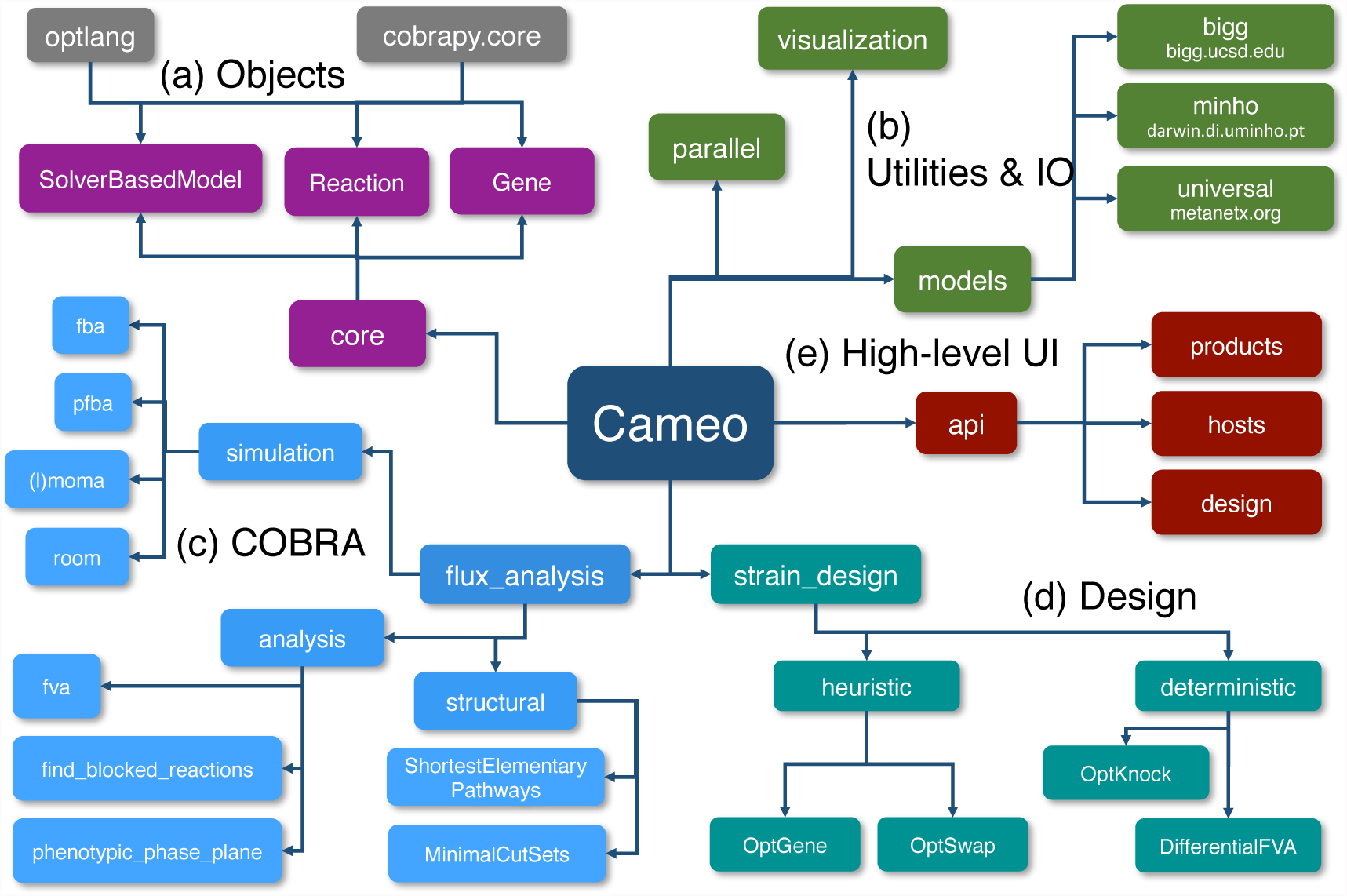
Package organization and functionality overview. The cameo package is organized into a number of sub-packages: *core* extends cobrapy’s own *core* package (Ebrahim et al., 2015) to use optlang (Jensen et al., 2017) as interface to a number of optimization solvers. *models* enables programmatic access to models hosted on the internet. *parallel* provides tools for the parallelization of design methods. *visualization* provides a number of high-level visualization functions, e.g., production envelopes. *flux analysis* implements many basic simulation and analysis methods needed for higher-level design methods and the evaluation of production goals etc. *strain design* provides a collection of *in silico* design methods and is subdivided into methods that use deterministic and heuristic optimization approaches. At last, *api* provides a high-level interface for computing designs.

To avoid duplication of effort, cameo is based on the constraint-based modeling tool cobrapy (Ebrahim et al., 2013) thus providing its users with already familiar objects and methods (see also Figure 2a). Furthermore, cameo takes advantage of other popular tools of the scientific Python stack, like for example Jupyter notebooks for providing an interactive modeling environment (Pérez and Granger, 2007) and pandas for the representation, querying, and visualization of results (McKinney, 2010).

Accessing published GEMs can be a challenging task as they are often made available in formats that are not supported by existing modeling software (Ebrahim et al., 2015). Cameo provides programmatic access to collections of models (Figure 2b) hosted by BiGG (King et al., 2016) and the University of Minho darwin.di.uminho.pt/models. Furthermore, by relying on the common namespace for reaction and metabolite identifiers provided by the MetaNetX.org project (Bernard et al., 2014) that covers commonly used pathway databases like KEGG (Kanehisa et al., 2016), RHEA (Morgat et al., 2015), and BRENDA (Chang et al., 2015), a universal reaction database can be used to predict heterologous pathways (see Supplementary Notebook 7 [v0.10.3, current].

Most design algorithms rely on solving optimization problems. In order to speed up simulations and ease the formulation of optimization problems, cameo replaces the solver interfaces utilized in cobrapy with optlang (Jensen et al., 2017), a Python interface to commonly used optimization solvers and symbolic modeling language that is maintained by the authors of cameo. Cameo always maintains a one-to-one correspondence of the GEM and its underlying optimization problem, greatly facilitating debugging and efficient solving by enabling warm starts from previously found solutions (Gelius-Dietrich et al., 2013). Furthermore, being based on sympy (SymPy Development Team, 2016), optlang enables the formulation of complicated optimization problems using symbolic math expressions, making the implementation of published design methods straightforward.

Runtimes of design methods are usually on the order of seconds to minutes. Nevertheless, scanning large numbers of potential products, host organisms, and feedstocks, can quickly make computations challenging (running the entire workflow using the high-level API takes on the order of hours). As described above, cameo makes unit operations as fast as possible by implementing an efficient interface to the underlying optimization software. In addition, a number of methods in cameo can be parallelized, and can thus take advantage of multicore CPUs and HPC infrastructure if available (see documentation).

With this broad overview of capabilities, we would like to emphasize the role of cameo as a useful resource to the modeling community and wish to support its development as a community effort in the long run. The majority of published strain design algorithms have not been experimentally validated (Machado and Herrgård, 2015) and we believe that their inaccessibility to users is a major factor for the lack of validation. With cameo we hope to counteract this problem by making these methods accessible to the entire metabolic engineering community and also providing a platform for modelers to implement and publish novel methods.

## CONCLUSIONS

With cameo version 0.10.3 we release a tool that is ready to be used in metabolic engineering projects. It is under active development and future work will include interfacing cameo with genome-editing tools to streamline the translation of computed strain designs into laboratory protocols, modeling of fermentation processes to get estimates on titers and productivities, and include pathway predictions based on retrobiosynthesis including hypothetical biochemical conversions (Campodonico et al., 2014).

## ACKNOWLEDGMENTS

We would like to thank Kai Zhuang, Miguel Campodonico and Sumesh Sukumura for providing valuable feedback and bug reports as early users of cameo. This project has received funding from the European Union’s Horizon 2020 research and innovation programme under grant agreement No 686070. Furthermore, we acknowledge financial support from the Novo Nordisk Foundation.

